## Supplementary material for "A network of coiled-coil and actin-like proteins controls the cellular organization of magnetosome organelles in deep-branching magnetotactic bacteria": Supplmental results, figures, and tables

**Extended Results:**

**Protein abundance during hydrogen-induced inhibition of magnetosome synthesis**

To investigate whether hydrogen affects magnetosome protein expression we compared the proteomes of RS-1 cells grown under hydrogen or nitrogen using liquid chromatography-mass spectrometry. Most magnetosome gene cluster (MGC) proteins showed similar levels, except Mad10, which was over 20 times more abundant in nitrogen (Supplemental Figure S11 A and B). However, Mad10 is not responsible for magnetosome inhibition under hydrogen conditions, as indicated by its magnetosome formation phenotype. In hydrogen-grown cells, hydrogenases, a response regulator, a signaling protein, and the ferric uptake regulator, Fur, were more abundant, while nitrogen-grown cells had higher levels of dehydrogenases and a carbon storage regulator (Supplemental Figure S11 C-F). These findings suggest potential molecular mechanisms for hydrogen's effect on magnetosome formation, warranting further investigation in the future.

**Architecture of Chain organization proteins in RS-1**

For each chain organization gene deleted in RS-1, predicted protein structures were modeled to visualize their structural features. Mad20, Mad25, and Mad26 all exhibited similar protein structures, characterized by long coiled-coil filament-like regions (Supplemental Figure S8 C-E). Similar features are seen in other bacterial species such as the coiled-coil protein, Crescentin, found in *Caulobacter crescentus*[1]. These coiled-coil domains are predicted to facilitate strong protein-protein interactions, which was supported by bacterial two-hybrid assays, where all three proteins interacted extensively with each other as well as with other chain organization proteins (Figure 6). Mad10 and Mad23, in contrast, have much shorter coiled-coil regions (Supplemental Figure S1 A), which is also reflected in their 3D structural models (Supplemental Figure S8 A-B). While Mad10 lacks additional predicted domains, previous studies suggest it contains a magnetite-binding region[2]. Mad23, however, possesses a distinctive HEAT repeat domain in addition to its small coiled-coil region (Supplemental Figure S1 A and Supplemental Figure S8 B-ii). HEAT domains are typically associated with protein-protein interactions, a hypothesis further supported by bacterial two-hybrid assays, where Mad23 showed strong interactions with Mad10 and Mad25 (Figure 6).

Both Mad28 and MamK are predicted to contain actin-like domains and share highly similar predicted protein structures (Supplemental Figure S1 A and Supplemental Figure S8 F-G). To further investigate their relationship, we performed both 3D structural and protein sequence alignments. While the sequence alignment revealed low similarity (Supplemental

Figure S8 J), the 3D structural alignment showed strong alignment between the two proteins (Supplemental Figure S8 H), suggesting that despite sequence divergence, their structural similarities may indicate conserved functions. However, our genetic studies and biomineralization time course analyses revealed distinct roles: MamK is essential for subchain formation, while Mad28 is required for proper chain localization along the positive curvature (Figure 5F-G). Additionally, the original *mad28* sequence from RS-1 in the NCBI database before 2019 lacked the first 114 nucleotides or 38 amino acids, shown in green in Supplemental Figure S8 H-J. However, this region is essential for restoring the wild-type phenotype in *mad28* complementation experiments (Supplemental Figure S6 F). Notably, this sequence is unique to Mad28 and does not align with MamK in the 3D structural comparison (Supplemental Figure S8 H). This specific *mad28* region is also present in Mad28 homologs from other deep-branching MTB, further supporting its functional significance. Supplemental Figure S8-I highlights this region in green in MYR-1, a deep-branching MTB from the *Nitrospirota* phylum.

**Extended data Figure:**

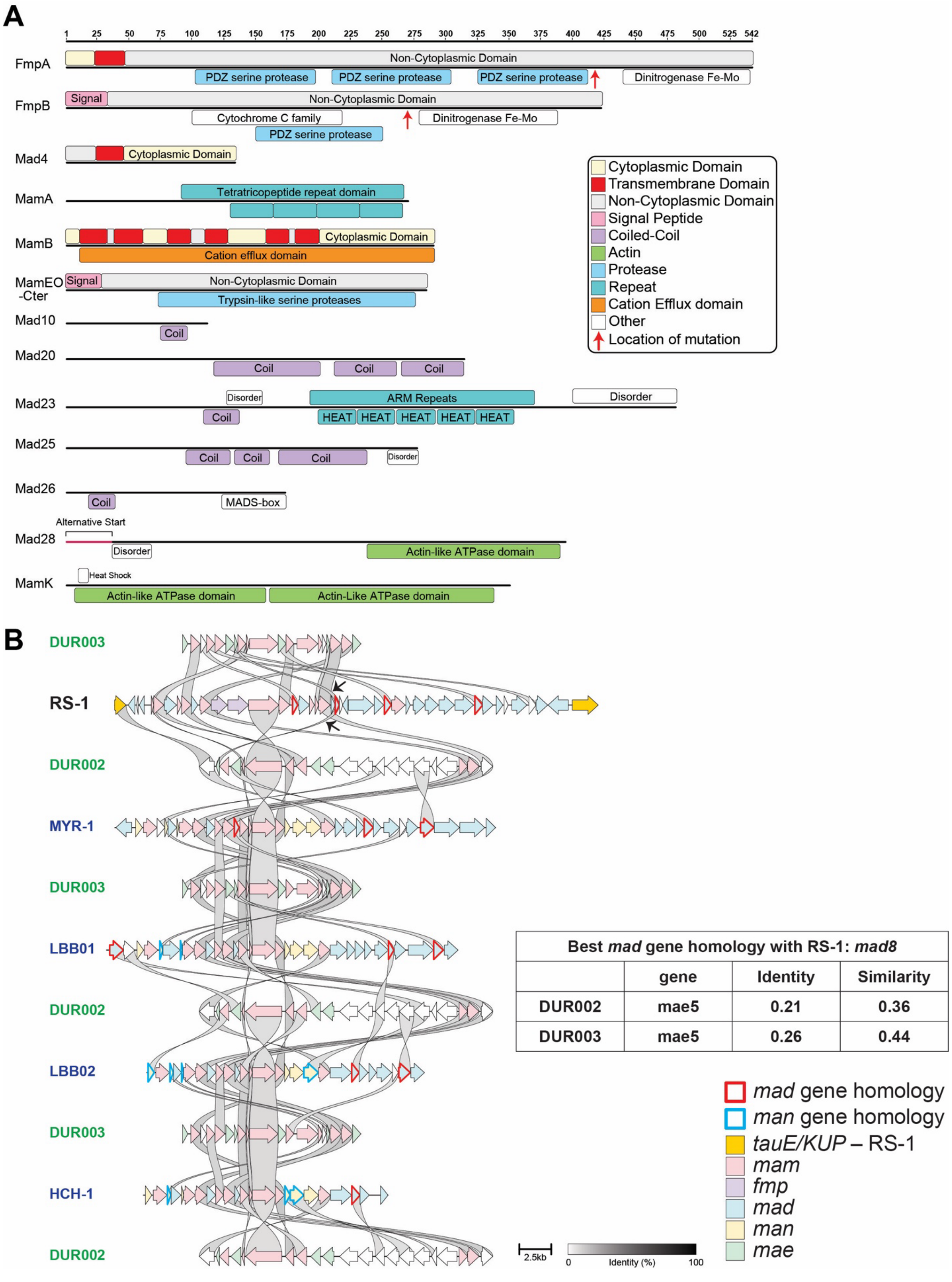

**Supplemental S1.** *Protein domains of magnetosome genes in this study and mad gene comparisons between deep branching MTB.*

(A) Domains of proteins studied here. Protein domains were identified using InterPro [3] and uploaded on to Geneious [4] to generate domain maps. Transmembrane domains were additionally verified using DeepTMHMM[5]. Red arrows on FmpA and FmpB indicate the location of their mutations for $fmpA^{Q418*}$ and $fmpB^{808delG}$ mutants. Mad28 shows an alternative start site that was not included in the original NCBI annotation. Importantly, Δ*mad28* mutant could only be complemented when the alternative start site was included. (B) Magnetosome gene cluster comparison between DUR002 and DUR003 *(Elusimicrobiota)* and RS-1 *(Desulfobacterota* representative*),* and between DUR002 and DUR003 and MYR-1, LBB01, LBB02, and HCH-1 *(Nitrospirota* representatives*)*. It has been stated that DUR002 and DUR003 do not contain any mad genes[6]. However, this analysis indicates that they do have homologues to some *mad* genes. MGCs were extracted from genomes available on NCBI manually and compared with Clinker[7]. Genes outlined in red are *mad* genes that show some homology in either DUR002 or DUR003. Genes outlined in blue are *man* genes that show some homology in either DUR002 or DUR003. The table on the right shows the proportions of identity and similarity between *mad8* of RS-1 and its homolog, *mae5*, which is found in both DUR002 and DUR003. The black arrows indicate the *mad8* homology on the comparison map. *mad4*, *mad11* and *mad24* from RS-1 also have homologs, but only in DUR003. In addition, *mad24* from HCH-1 and LBB02 has a homolog to a gene in the MGC of DUR003.

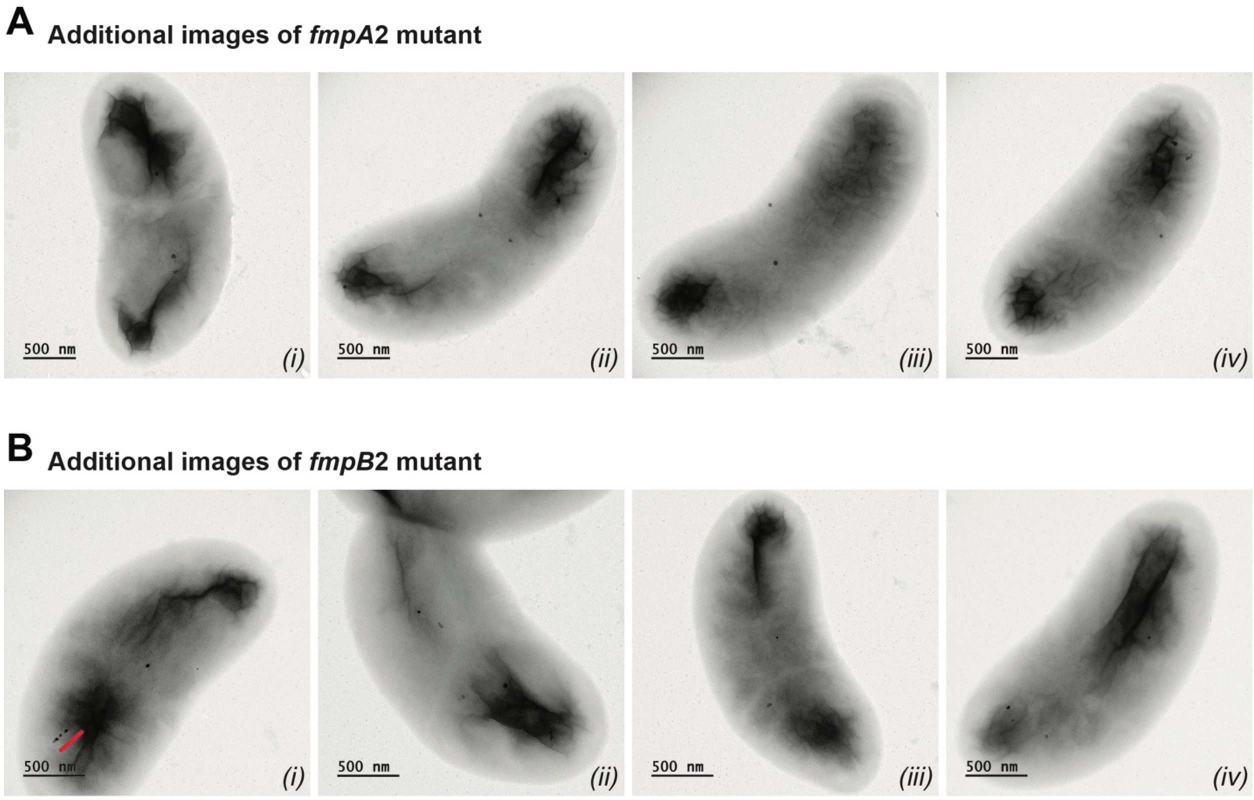


**Supplemental S2.** *Additional TEM images of fmpA and fmpB mutants.*

(A) Additional TEM images of *fmpA* mutant in RS-1. These images illustrate fewer and misshaped crystals in each cell to complement the statistical data from Figure 4 and Supplemental Figure S7. (B) Additional TEM images of *fmpB* mutant in RS-1. These images illustrate the fewer and smaller crystals in each cell to complement the statistical data from Figure 4 and Supplemental Figure S7. Additionally, B-*i* shows an example of a chain phenotype in this mutant.

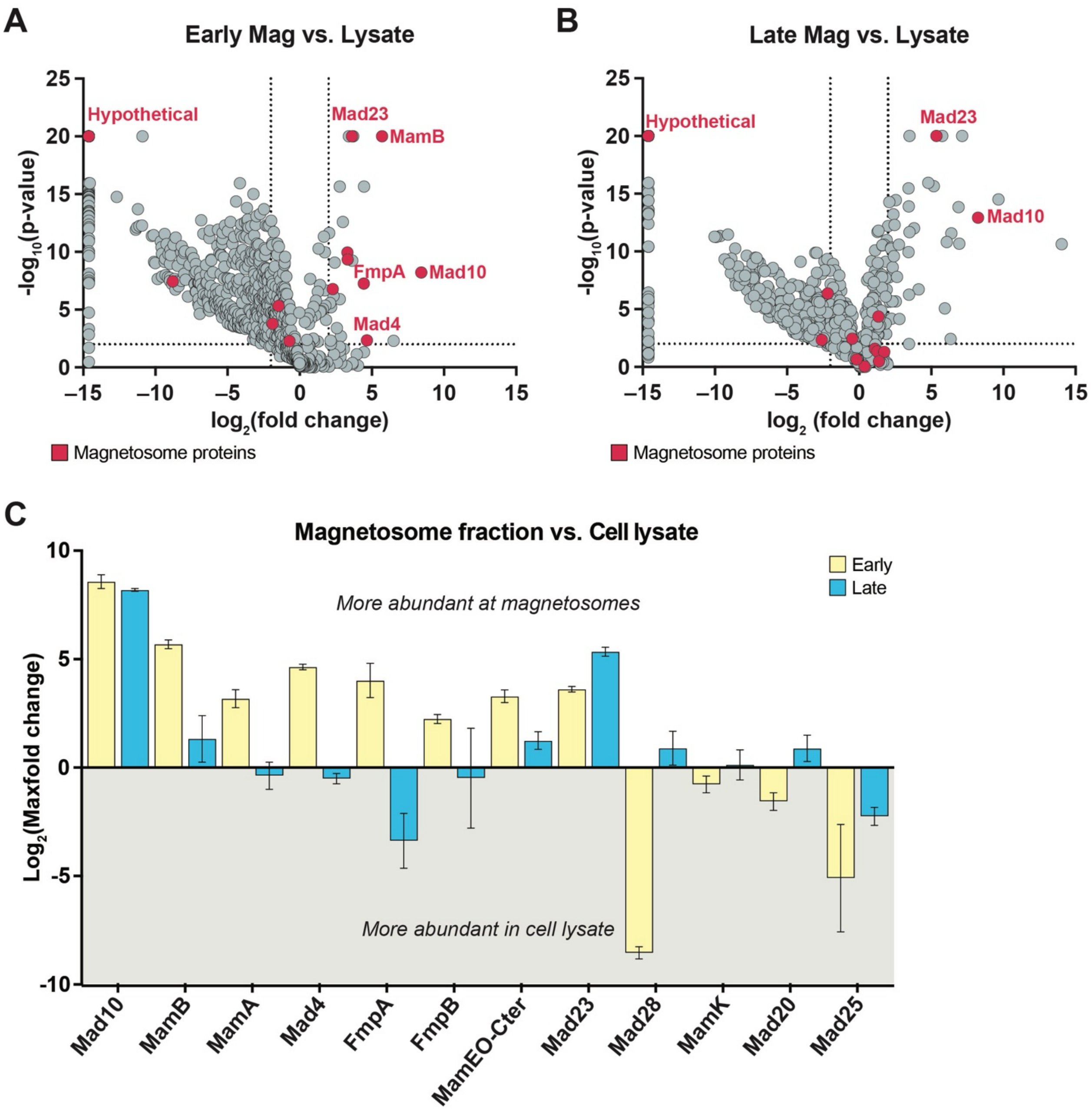


**Supplemental S3.** *Additional proteomics data for early vs. late stage of biomineralization.* (A-B) A volcano plots showing the differential abundance analysis between magnetosome fraction and whole cell lysate for the early (A) and late (B) stages of biomineralization. Red dots indicate proteins from the magnetosome gene cluster (DMR_040710 – DMR_41530). Proteins with a peptide abundance of 0 in the magnetosome fraction result in an infinite fold change. Therefore, their fold change was arbitrarily set to 25,000. This is reflected in the volcano plots as a $\log_2$ value of -15. Additionally, any p-value of 0 was arbitrarily set to a $-\log_{10}$(p-value) of 20. (C) Abundance of magnetosomes proteins (those encoded by MGC genes) in the cell lysate versus the magnetosome fraction. The abundance of magnetosome proteins is represented as greater than 0 when they are more prevalent in the magnetosome fraction and as less than 0 when they are more abundant in the cell lysate.

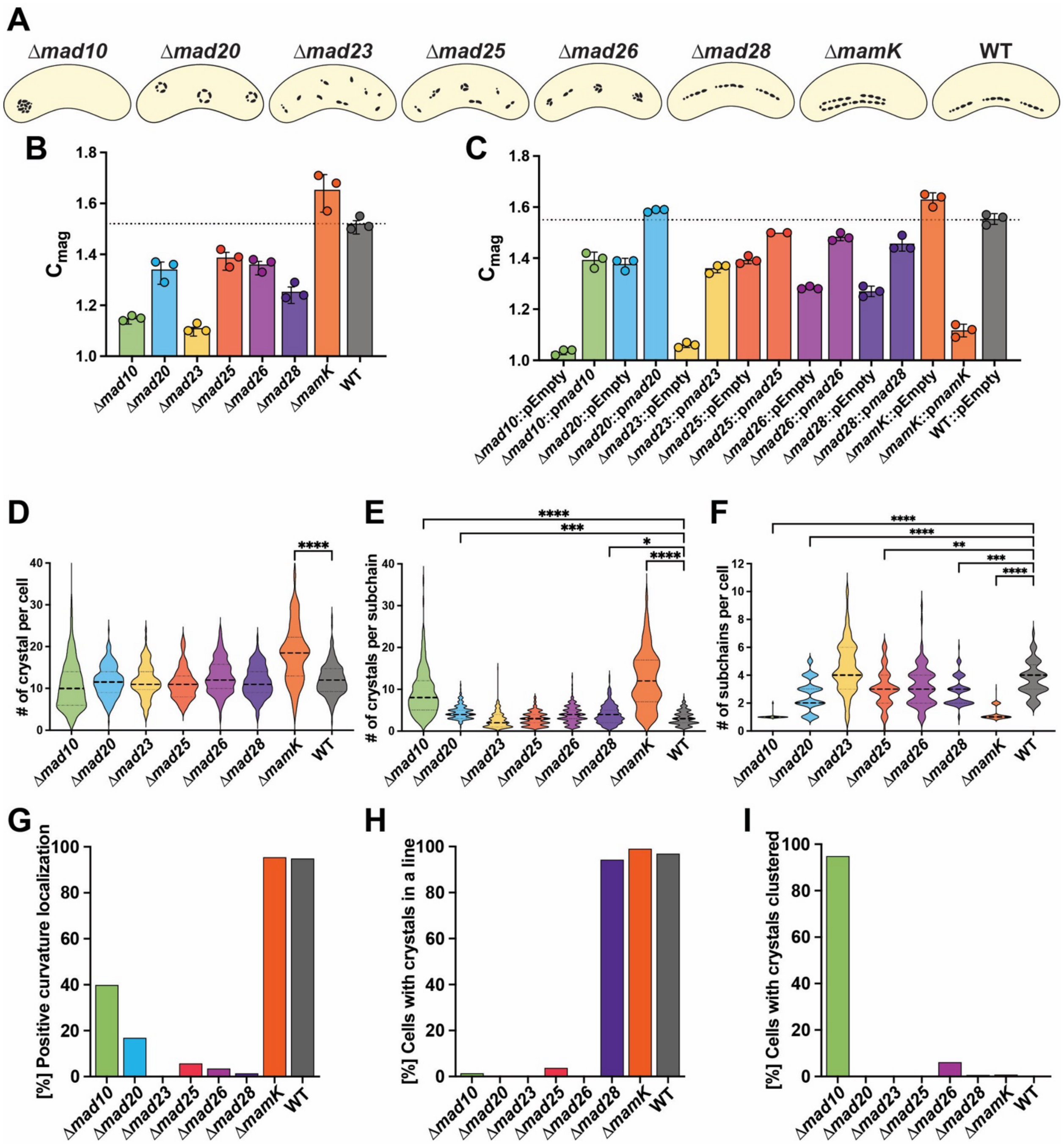


**Supplemental S4.** *Characterization of magnetosome gene deletions in RS-1*

(A) Cartoon diagram of representative phenotype of each strain. (B) $C_{Mag}$ values for each strain without plasmids. Error bars indicate standard deviation from average values of three cultures.

(C) $C_{Mag}$ values for each mutant with an empty plasmid or with a plasmid constitutively expressing the deleted gene. The WT strain carries only an empty plasmid. Error bars indicate standard deviation from average values of three cultures. The dotted line in (B) and (C) represents the average $C_{Mag}$ of WT. (D) Violin plots displaying the number of crystals per cell for each strain. (E) Violin plots illustrating the number of crystals per subchain for each strain. (F) Violin plots representing the number of subchains per cell for each strain. All statistical tests used on (D), (E), and (F) are stated in *Supplemental table 8*. (G-I) The percentage of cells exhibiting specific phenotypes, based on a count of 200 cells per strain. (G) The percentage of cells with crystals localized at the positive curvature of the cell. (H) The percentage of cells with magnetosomes displaying a chain phenotype. (I) The percentage of cells with all magnetosomes in the cell clustered together.

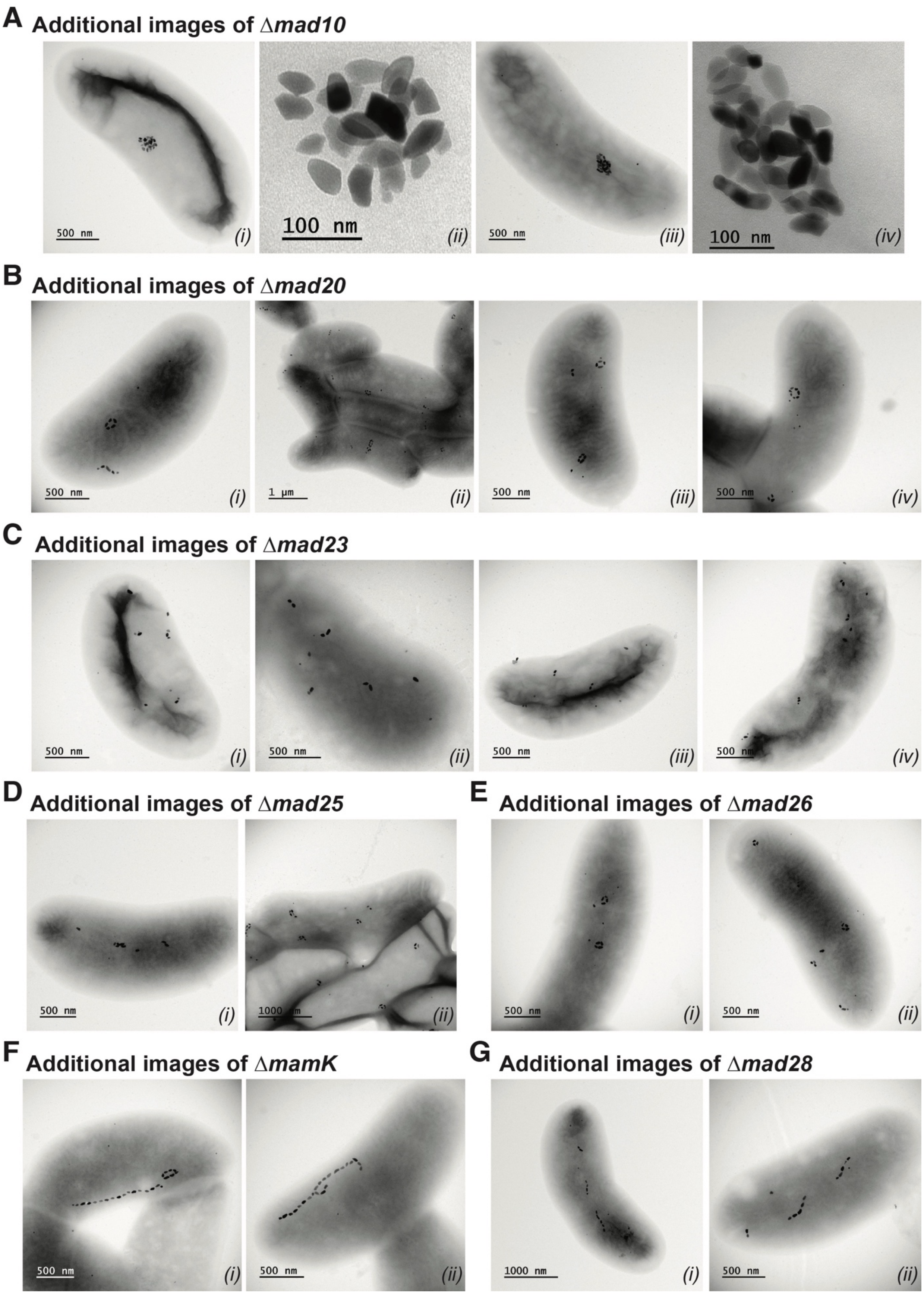
A Additional images of Δmad10
500 nm
(i)
100 nm
(ii)
500 nm
(iii)
100 nm
(iv)
B Additional images of Δmad20
500 nm
(i)
1 µm
(ii)
500 nm
(iii)
500 nm
(iv)
C Additional images of Δmad23
500 nm
(i)
500 nm
(ii)
500 nm
(iii)
500 nm
(iv)
D Additional images of Δmad25
500 nm
(i)
1000 nm
(ii)
E Additional images of Δmad26
500 nm
(i)
500 nm
(ii)
F Additional images of ΔmamK
500 nm
(i)
500 nm
(ii)
G Additional images of Δmad28
1000 nm
(i)
500 nm
(ii)

111

**Supplemental S5.** *Additional TEM images of gene deletions in RS-1 from this study.* (A) Additional TEM images of Δ*mad10* mutant. (A-*ii* and *iv*) are zoomed in images of the cluster of magnetosomes from A-*i* and *iii* respectively and illustrate the overlapping of magnetosomes. (B) Additional TEM images of Δ*mad20* mutant. These images illustrate fewer subchains in each cell and more crystals per subchain in each cell to complement the statistical data from Supplemental Figure S1. Perhaps rings form in this mutant because subchains are too long, similar to the Δ*mamK* phenotype. (C) Additional TEM images of Δ*mad23* in RS-1. These images illustrate the higher number of subchains in each cell and fewer crystals per subchain in each cell to complement the statistical data from Supplemental Figure S1. Subchains are more scattered in Δ*mad23*. (D) Additional TEM images of Δ*mad25* in RS-1. These images are to complement the statistical data from Supplemental Figure S1. (E) Additional TEM images of Δ*mad26* in RS-1. These images are to complement the statistical data from Supplemental Figure S1. Subchains are clustered in Δ*mad25* and Δ*mad26*. (F) Additional TEM images of Δ*mamK* in RS-1. These images illustrate the higher number of crystals in each cell and absence of subchains to complement the statistical data from Supplemental Figure S1. Additionally, the placement of the chain is not always in the center of the cell as seen in F-*ii*. (G) Additional TEM images of Δ*mad28* in RS-1. These images illustrate that chain placement is not at the positive curvature of the cell and to complement the statistical data from Supplemental Figure S1.

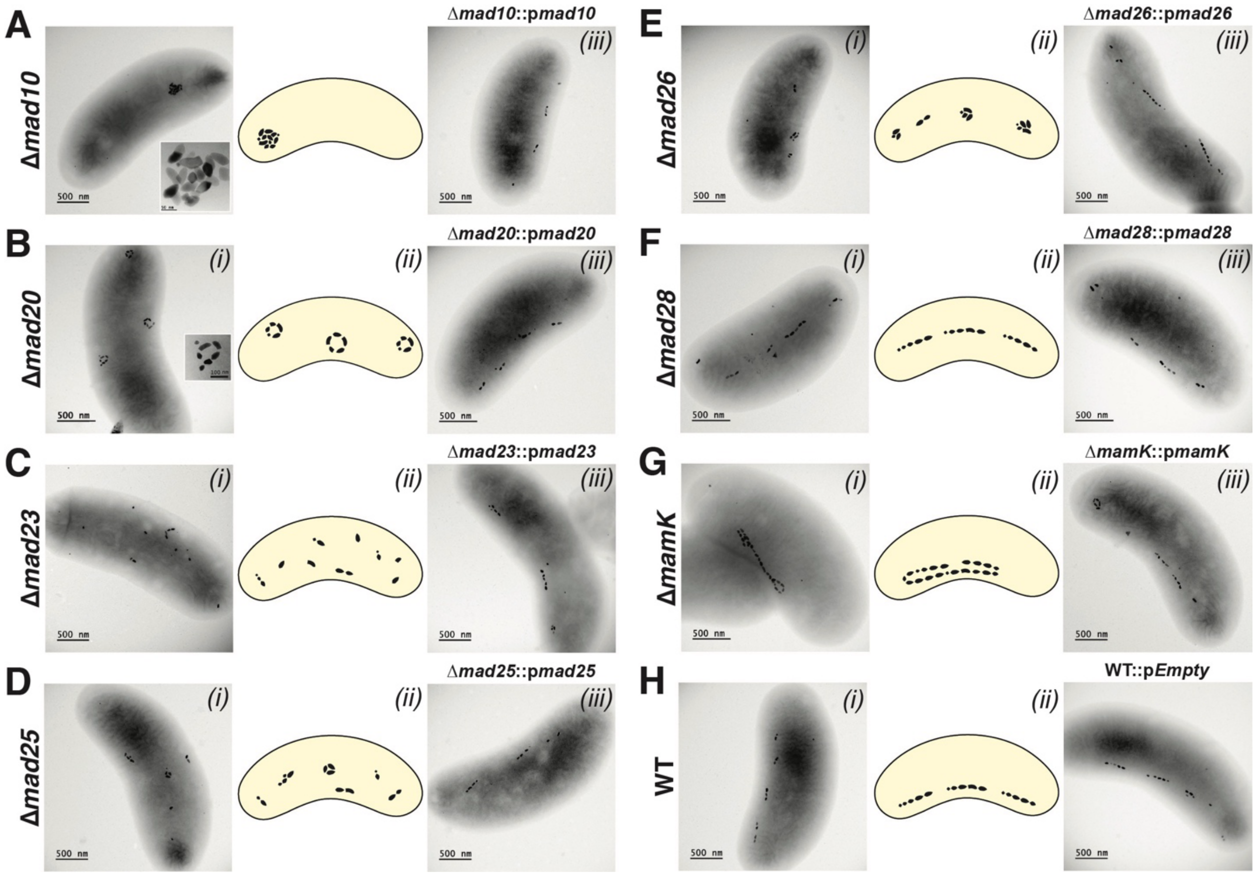


**Supplemental S6.** *TEM images of complementations of deletion mutants*

(A-G) TEM images of deletion mutants and their respective complementation strains in RS-1. All complementations were done by expressing the gene on a plasmid. (H) TEM images of WT and a WT control with an empty plasmid.

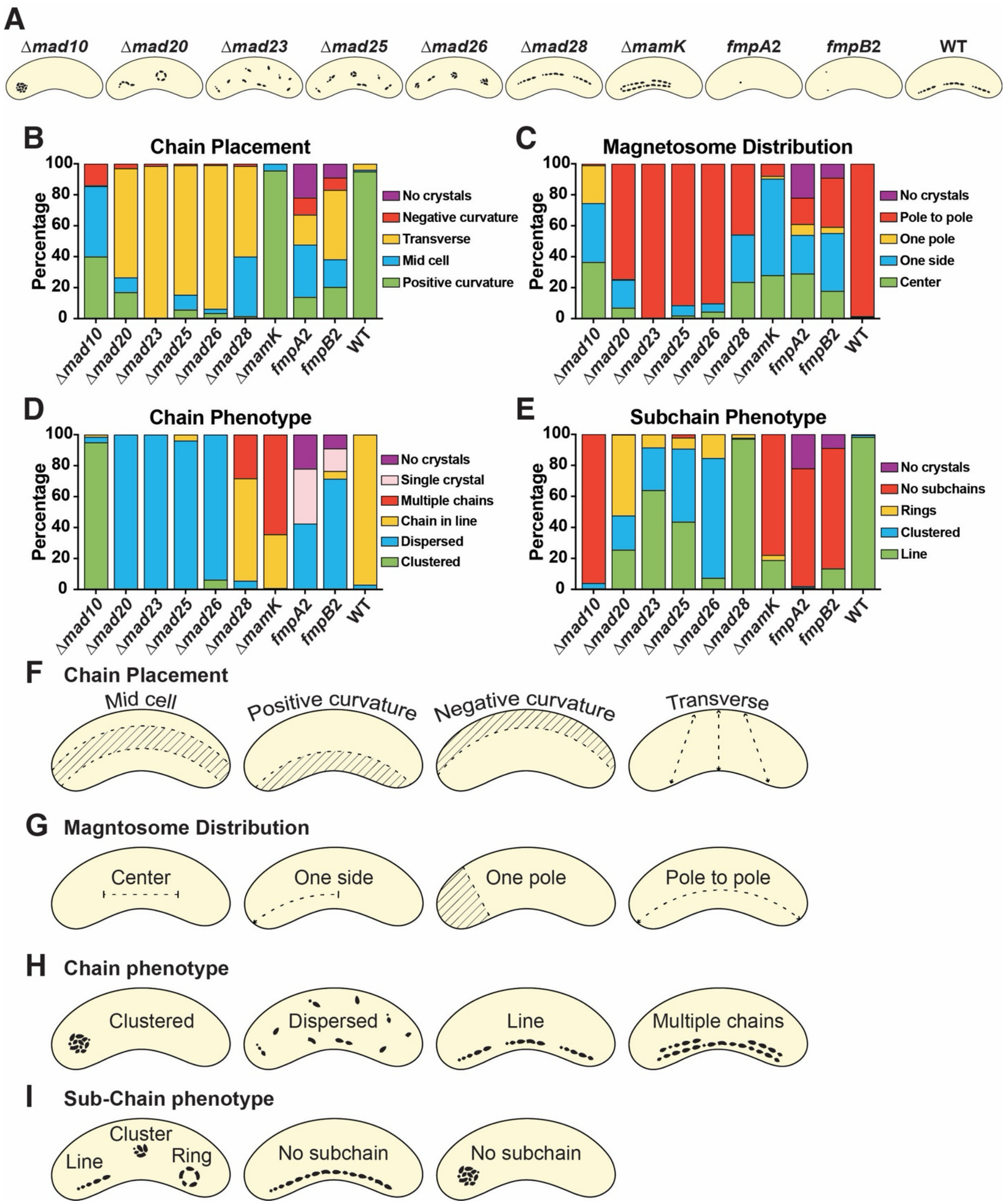


**Supplemental S7.** *Additional mutant phenotypes and magnetosome chain and subchain categories*

(A) Illustrations of the phenotypes of the different mutants in this study. (B-E) Additional phenotype categories for all mutants. Over 200 cells were counted for each mutant. (F-I)

Categories used to calculate mutant phenotypes in Figure 4, Supplemental Figure S4, as well as (B-E) in this figure.

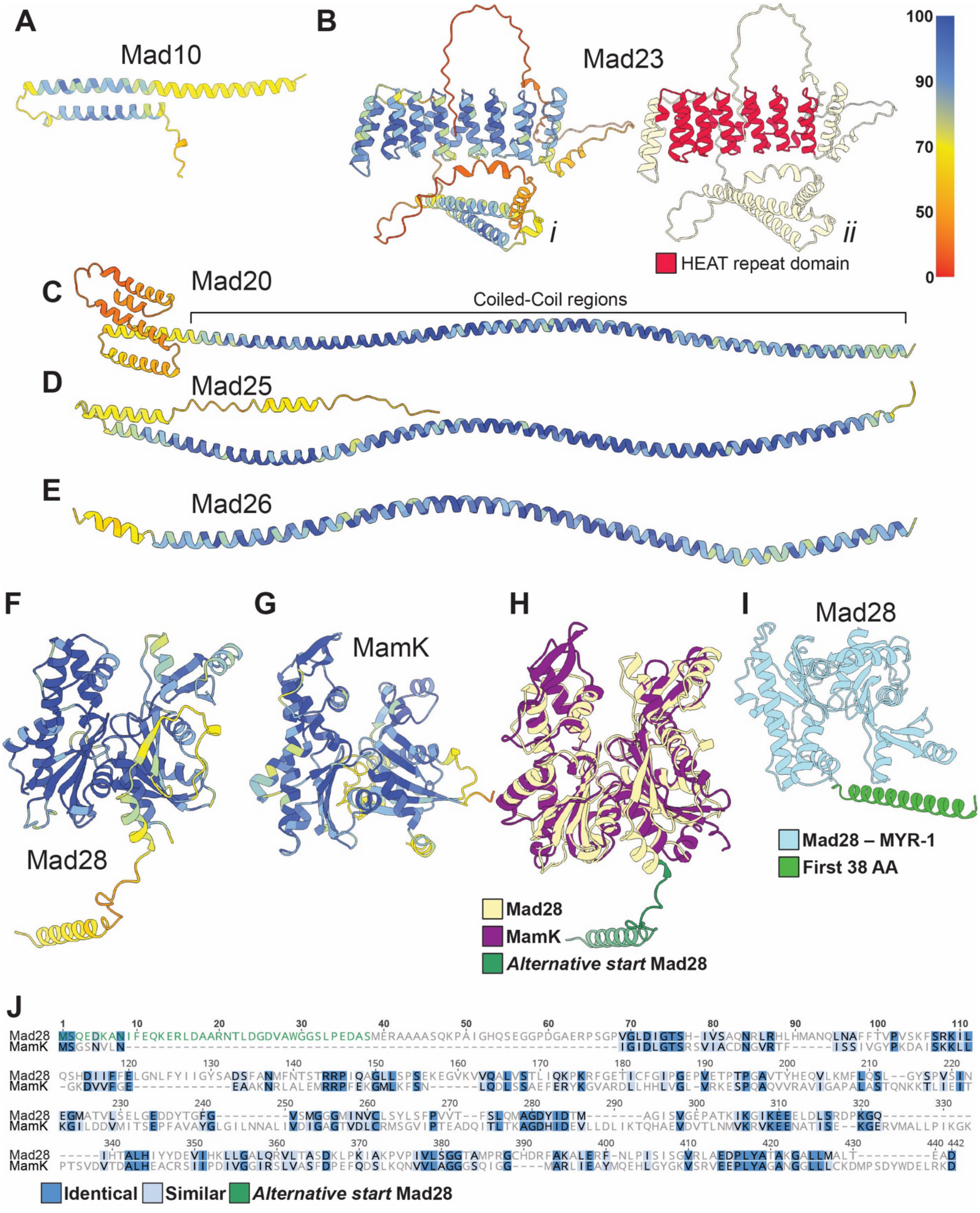

**Supplemental S8.** *Predicted 3D Structures of magnetosome proteins deleted in RS-1 and alignments of MamK and Mad28*

(A-I) Predicted protein structures of Mad proteins and MamK in this study were made with Alphafold3[8] and visualized with ChimeraX[9]. 3D structures are colored according to Alphafold confidence values, with blue at 100% confident and red 0% confidence. (B-*ii*) Indicates the location of the HEAT domain (colored in red) in Mad23. (H) 3D alignment of protein structures of Mad28 and MamK from RS-1. Protein structures were made with alphafold3 and then a 3D alignment was preformed using ChimeraX[9]. Purple is the 3D predicted structure for MamK, yellow is the predicted structure for Mad28 and green is the part of the structure produced from the *mad28* alternative start codon. (I) Predicted 3D structure of Mad28 from MYR-1, a deep-branching MTB in the *Nitrospirota* phylum. The first 38 amino acids are highlighted in green to indicate structural similarity to the corresponding region in RS-1's Mad28. (J) 2D protein sequence alignment of MamK and Mad28 using MAFFT v7.490 plug-in on Geneious[4]. Blue highlighted residues show similarities between sequences and dashes represent gaps. Green letters show the sequence for the alternative start in Mad28.

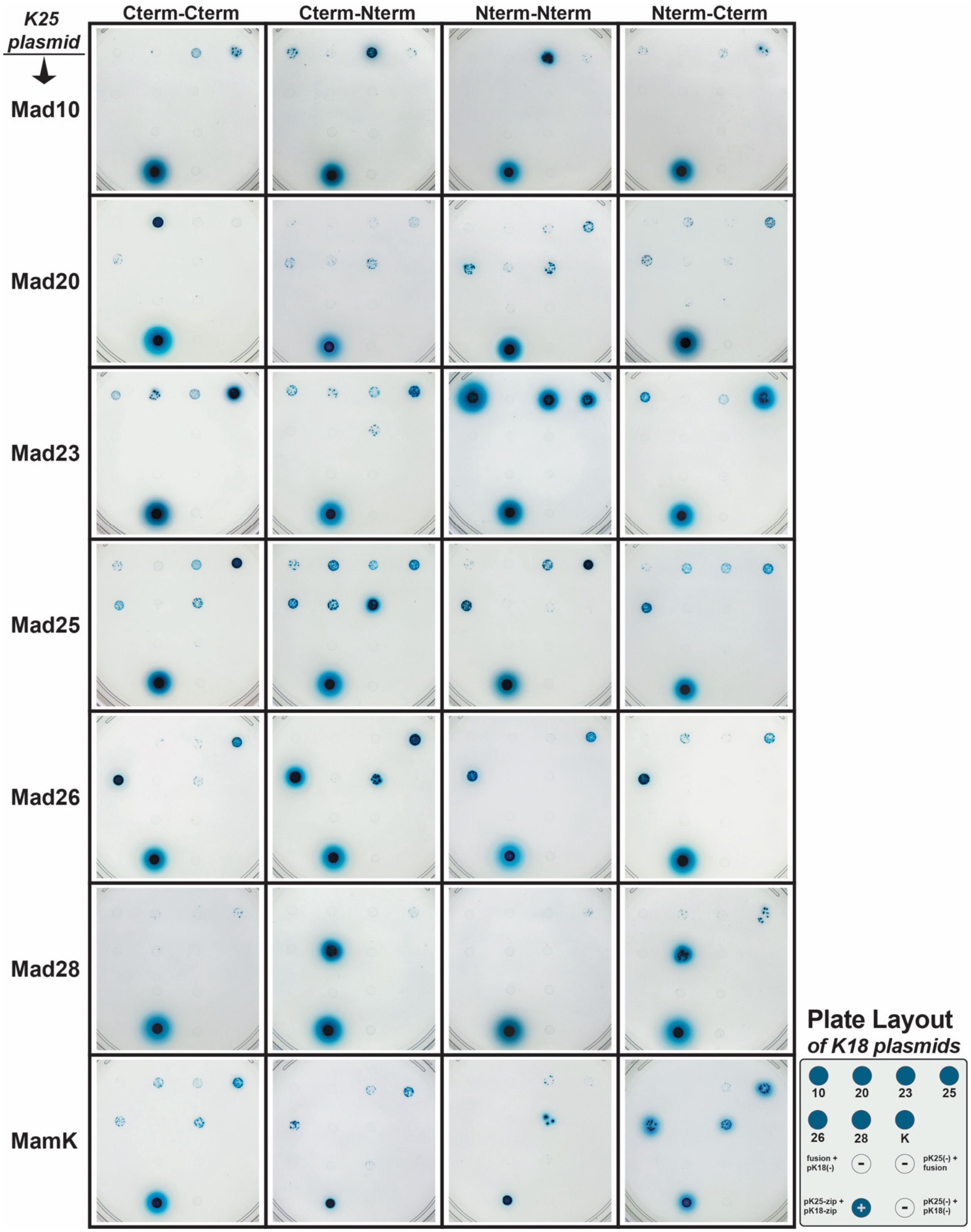


162
163 **Supplemental S9.** *BACTH images of plates for all interactions.*

Image of all plates used to construct the table of the interactions in Figure 6. The illustration on the bottom right shows the layout for each plate and the controls used.

**A**

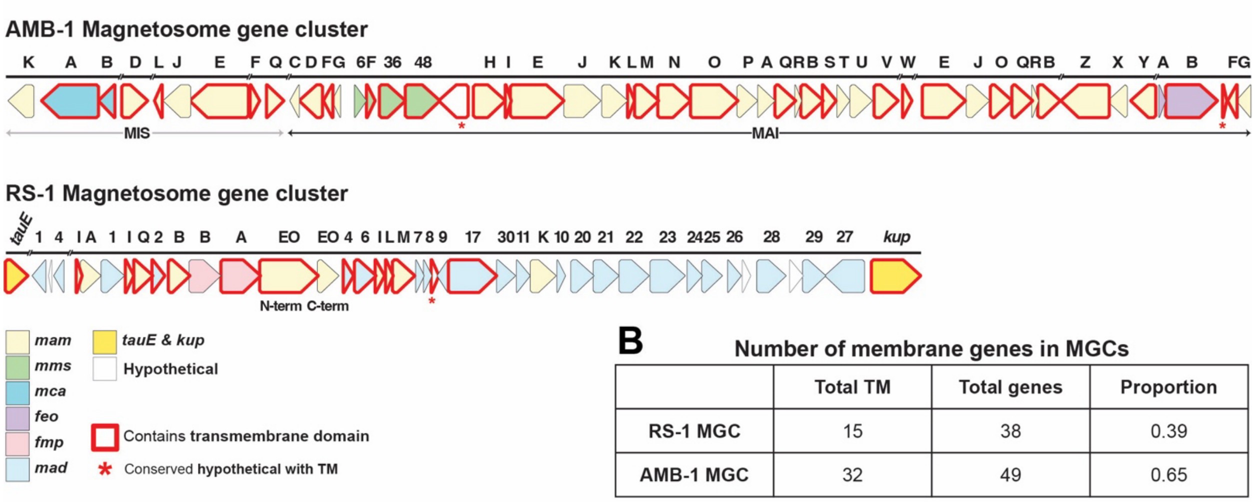


**B** Number of membrane genes in MGCs

| | Total TM | Total genes | Proportion |
|---|---|---|---|
| RS-1 MGC | 15 | 38 | 0.39 |
| AMB-1 MGC | 32 | 49 | 0.65 |

**C**

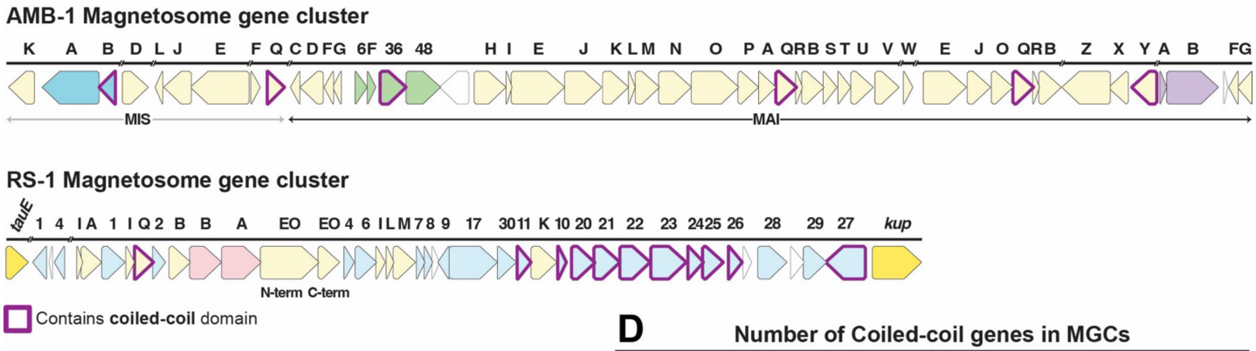


**D** Number of Coiled-coil genes in MGCs

| | Total CC | Total genes | Proportion |
|---|---|---|---|
| RS-1 MGC | 10 | 38 | 0.26 |
| AMB-1 MGC | 6 | 49 | 0.12 |

**Supplemental S10.** *Magnetosome gene cluster comparison between AMB-1 and RS-1.* (A) Magnetosome gene clusters in AMB-1 and RS-1. Genes highlighted in red encode proteins with predicted transmembrane domains. (B) Table of the proportions of proteins with predicted transmembrane domains to all proteins encoded by the magnetosome gene clusters of AMB-1 and RS-1. DeepTMHMM[5] was used to gather transmembrane domain predictions. Hypothetical genes were not considered in the calculations for the proportions. (C) Magnetosome gene cluster of AMB-1 and RS-1 with genes encoding proteins with predicted coiled-coil domains highlighted in purple. (D) Table of the proportions of proteins with coiled-coil domains compared to all proteins in the MGC for AMB-1 and RS-1. DeepCoil[10] was used to obtain coiled-coil predictions. A threshold of 0.35 or greater significance with positive predictions for *a* and *d* core positions was used to verify coiled-coil predictions. Hypothetical genes also were not considered in the calculations for the proportions of coiled-coil genes.

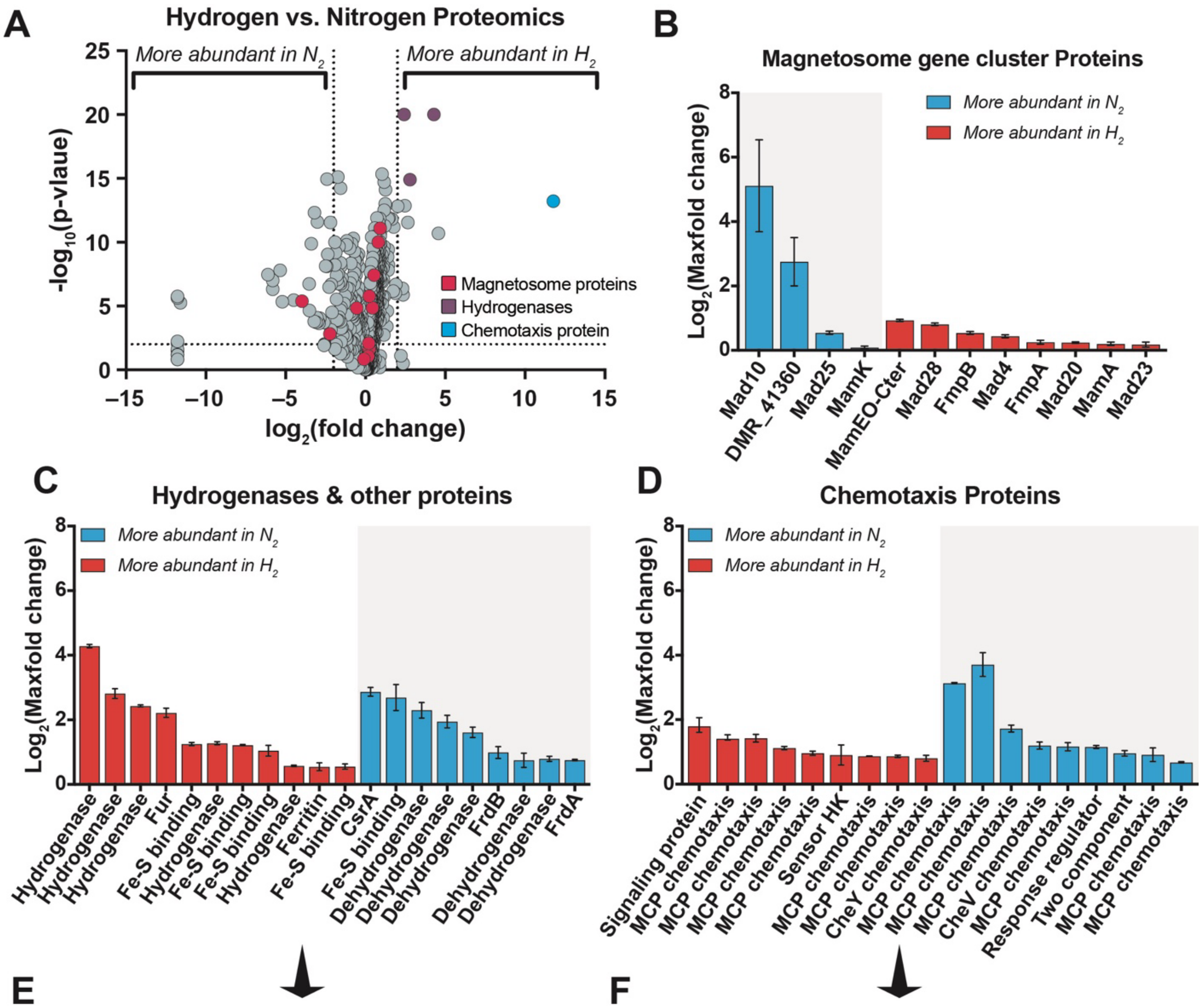


**E**

| | Protein name | Highest |
|---|---|---|
| 1 | DMR_07840 Hydrogenase | Hydrogen |
| 2 | DMR_07830 Hydrogenase | Hydrogen |
| 3 | DMR_07850 Hydrogenase | Hydrogen |
| 4 | DMR_19790 Fur ferric uptake regulation | Hydrogen |
| 5 | DMR_01950 Fe-S binding | Hydrogen |
| 6 | DMR_01930 Hydrogenase | Hydrogen |
| 7 | DMR_01940 Fe-S binding | Hydrogen |
| 8 | DMR_20050 Fe-S binding | Hydrogen |
| 9 | DMR_12950 Hydrogenase | Hydrogen |
| 10 | DMR_17620 Ferritin | Hydrogen |
| 11 | DMR_42710 Fe-S binding | Hydrogen |
| 12 | DMR_31480 CsrA carbon storage regulator | Nitrogen |
| 13 | DMR_42500 Fe-S binding | Nitrogen |
| 14 | DMR_01890 Dehydrogenase | Nitrogen |
| 15 | DMR_16050 Dehydrogenase | Nitrogen |
| 16 | DMR_21490 Dehydrogenase | Nitrogen |
| 17 | DMR_05780 FrdB fumarate reductase | Nitrogen |
| 18 | DMR_19380 Dehydrogenase | Nitrogen |
| 19 | DMR_08200 Dehydrogenase | Nitrogen |
| 20 | DMR_34280 FrdA fumarate reductase | Nitrogen |

**F**

| | Protein name | Highest |
|---|---|---|
| 1 | DMR_09360 Signaling protein | Hydrogen |
| 2 | DMR_28950 Methyl-accepting chemotaxis | Hydrogen |
| 3 | DMR_00340 Methyl-accepting chemotaxis | Hydrogen |
| 4 | DMR_37860 Methyl-accepting chemotaxis | Hydrogen |
| 5 | DMR_27200 Methyl-accepting chemotaxis | Hydrogen |
| 6 | DMR_34160 Sensor histidine kinase | Hydrogen |
| 7 | DMR_38310 Methyl-accepting chemotaxis | Hydrogen |
| 8 | DMR_04320 Methyl-accepting chemotaxis | Hydrogen |
| 9 | DMR_23920 CheY chemotaxis | Hydrogen |
| 10 | DMR_26500 Methyl-accepting chemotaxis | Nitrogen |
| 11 | DMR_07950 Methyl-accepting chemotaxis | Nitrogen |
| 12 | DMR_29030 Methyl-accepting chemotaxis | Nitrogen |
| 13 | DMR_26640 CheV chemotaxis | Nitrogen |
| 14 | DMR_07730 Methyl-accepting chemotaxis | Nitrogen |
| 15 | DMR_41540 Response regulator | Nitrogen |
| 16 | DMR_45100 2-component sensor & regulator | Nitrogen |
| 17 | DMR_08750 Methyl-accepting chemotaxis | Nitrogen |
| 18 | DMR_27400 Methyl-accepting chemotaxis | Nitrogen |

**Supplemental S11.** *Additional proteomics data for hydrogen vs nitrogen.*

(A) A volcano plot showing the differential abundance analysis of the proteome between hydrogen and nitrogen conditions. Red dots indicate magnetosome genes, most of which fall between -2 and 2 $\log_2$ fold change, indicating no significant difference in abundance. Purple dots indicate hydrogenase proteins and blue dot indicates a chemotaxis protein, all of which are greater than 2 $\log_2$ fold more abundant in the hydrogen condition. Proteins with a peptide abundance of 0 in one condition may result in an infinite fold change. In such cases, the fold change was arbitrarily set to 3500. This is reflected in the volcano plots as a $\log_2$ value of -11.8. Additionally, p-values of 0 were arbitrarily assigned a $-\log_{10}$(p-value) of 20. (B) The maximum fold change of magnetosome proteins detected in cell lysates under both nitrogen and hydrogen conditions. Blue bars represent proteins more abundant in nitrogen condition and orange bars represent proteins more abundant in hydrogen condition. (C) and (D) Additional proteomic data comparing protein abundance in nitrogen versus hydrogen conditions, highlighting non-magnetosome proteins with maximum fold changes above 2. (E) and (F) display the gene names and locus IDs corresponding to all bars in (C) and (D), respectively.

**Extended data Table:**

| Phyla | Organiam | 16S gene ID | Genome ID | Shape reference | MGC reference |
|---|---|---|---|---|---|
| *Alpha* | *Magnetospirillum magneticum* AMB-1 | D17514.1 | NCBI:txid342108 | [11] | [12] |
| *Alpha* | *Magnetospirillum magnetotacticum* MS-1 | IMG: 2647169949 | NCBI:txid272627 | [13] | [14] |
| *Alpha* | *Magnetospirillum caucaseum* SO-1 | JX502622.2 | NCBI:txid1244869 | [15] | [15–17] |
| *Alpha* | *Magnetospirillum* ME-1 | IMG: 2676550241 | NCBI:txid1639348 | [18] | [16,17,19] |
| *Alpha* | *Magnetospirillum* XM-1 | KP966105.1 | NCBI:txid1663591 | [20] | [16,21] |
| *Alpha* | *Magnetospirillum marisnigri* SP-1 | KC252630.3 | NCBI:txid1285242 | [15] | [17,22] |
| *Alpha* | *Magnetospirillum kuznetsovii* LBB-42 | NR_180134.1 | NCBI:txid2053833 | [23] | [16,23] |
| *Alpha* | *Magnetospirillum gryphiswaldense* MSR-1 | NR_121771.1 | NCBI:txid431944 | [24,25] | [26] |
| *Alpha* | *Magnetospirillum moscoviense* BB-1 | KF712468.2 | NCBI:txid1437059 | [15] | [17,22] |
| *Alpha* | *Magnetospirillum* sp. LM-5 | IMG: 2894467686 | NCBI:txid2681466 | [27] | [17] |
| *Alpha* | *Magnetospirillum* sp. SS-4 | IMG: 2894461381 | NCBI:txid2681465 | [27] | [17] |
| *Alpha* | *Magnetospirillum* sp. UT-4 | IMG: 2894472958 | NCBI:txid2681467 | [27] | [17] |
| *Alpha* | *Ca. Magneticavibrio boulderlitore* LM-1 | JF490044.1 | NCBI:txid1008128 | [27] | [17] |
| *Alpha* | *Magnetospira* QH-2 | EU675666.1 | NCBI:txid1288970 | [28] | [29] |

| | | | | | |
|---|---|---|---|---|---|
| *Alpha* | *Ca. Terasakiella magnetica* PR1 | IMG: 2752957094 | NCBI:txid1867952 | [30] | [30] |
| *Alpha* | *Magnetovibrio blakemorei* MV-1 | L06455.1 | NCBI:txid28181 | [31] | [32] |
| *Gamma* | Strain BW-2 | HQ595728.1 | NCBI:txid947515 | [33] | [34] |
| *Gamma* | SHHR-1 | KX344069.1 | NCBI:txid1899433 | [35] | [16] |
| *Gamma* | Strain SS-5 | HQ595729.1 | NCBI:txid947516 | [33] | [36] |
| *Eta* | *Ca. Magnetococcus massalia* MO-1 | EF643520.2 | NCBI:txid451514 | [37] | [38] |
| *Eta* | *Magnetococcus* MC-1 | IMG: 640720050 | NCBI:txid156889 | [39] | [40] |
| *Eta* | *Magnetofaba australis* IT-1 | JX534168.1 | NCBI:txid1434232 | [41] | [41] |
| *Eta* | *Ca. Magnetaquicoccus inordinatus* UR-1 | IMG: 2881921666 | NCBI:txid2496818 | [42] | [42] |
| *Desulfo-bacterota* | *Ca. Magnetoglobus multicellularis* | IMG: 2559591426 | NCBI:txid890399 | [43,44] | [45] |
| *Desulfo-bacterota* | *Ca. Magnetananas rongchenensis* RPA | KF925363.1 | NCBI:txid1463558 | [46] | [46] |
| *Desulfo-bacterota* | *Ca. Magnetomorum* HK-1 | IMG: 2648890082 | NCBI:txid1509431 | [47] | [48] |
| *Desulfo-bacterota* | *Desulfamplus magnetovallimortis* BW-1 | JN252194.1 | NCBI:txid1073250 | [49] | [50] |
| *Desulfo-bacterota* | *Desulfovibrio magneticus* RS-1 | NR_074958.1 | NCBI:txid573370 | [51,52] | [53] |
| *Desulfo-bacterota* | *Desulfovibrio* FSS-1 | LC311577.1 | NCBI:txid2730080 | [54] | [54] |
| *Desulfo-bacterota* | *Delta Proteobacterium* FH-1 | JF330268 | NCBI:txid999117 | [50] | [50] |

| | | | | | |
|---|---|---|---|---|---|
| *Desulfo-bacterota* | *Desulfonatronum* ML-1 | HQ595725.1 | NCBI:txid947513 | 55 | 50 |
| *Nitro-spirota* | *Ca. Magnetoovum chiemensis* WYHC-5 | OL423397.1 | NCBI:txid1609970 | 56 | 56 |
| *Nitro-spirota* | *Ca. Magnetobacterium casensis* MYR-1 | MT703955.1 | NCBI:txid1455061 | 57 | 58 |
| *Nitro-spirota* | *Ca. Magnetobacterium bavaricum* TM-1 | X71838.1 | NCBI:txid29290 | 59 | 60 |
| *Nitro-spirota* | *Ca. Nitrospirae* YQR-1 | ON340538.1 | | 16 | 16 |
| *Nitro-spirota* | *Ca. Magnetomonas plexicatena* LBB-01 | MK632185.1 | CP049016.1 | 61 | 61 |
| *Nitro-spirota* | *Ca. Magnetominusculus linsii* LBB-02 | MK632186.1 | | 61 | 61 |
| *Nitro-spirota* | *Ca. Nitrospirae* MYC-10 | ON342894.1 | | 16 | 16 |
| *Nitro-spirota* | *Ca. Magnetocorallium paracelense* XS-1 | OQ281288.1 | NCBI:txid3021403 | 62 | 62 |
| *Elusi-microbiota* | *Ca. Liberimonas magnetica DUR002* | KY516811 | GCA_020523885.1 | 6 | 6 |
| *Elusi-microbiota* | *Ca. Obscuribacterium magneticum DUR003* | JQ369221 | GCA_020523905.1 | 6 | 6 |
| *OP3* | *Ca. Omnitrophus magneticus* SKK-01 | JN412733.1 | NCBI:txid174292 | 63 | 60 |

**Supplemental Table 1.** *Table of species and reference used for phylogenetic tree construction*

**Supplemental Tables**

| **Strain** | **Description** | **Reference** |
|---|---|---|
| AK80 | WT | [64] |
| AK180 | *fmpA2*$^{Q418*}$ | [64] |
| AK182 | *fmpB2*$^{808delG}$ | [64] |
| AK268 | WT, Δ*upp* | [65] |
| AK350 | Δ*mad23*, Δ*upp* | This study |
| AK377 | Δ*mad20*, Δ*upp* | This study |
| AK378 | Δ*mad25*, Δ*upp* | This study |
| AK380 | Δ*mad10*, Δ*upp* | This study |
| AK382 | Δ*mad28*, Δ*upp* | This study |
| AK383 | Δ*mad26*, Δ*upp* | This study |
| AK220 | Δ*mamK* | This study |

**Supplemental Table 2.** *Table of all strains used in this study*

| **Plasmid** | **Description** | **Backbone** | **Reference** |
|---|---|---|---|
| PAK906 | Complementation backbone with pNTP | | [64] |
| PAK907 | Complementation backbone with pmamA | | [64] |
| pAK914 | Deletion backbone with SacB only | | [64] |
| pAK1127 | Deletion backbone with Upp and SacB | | [65] |
| pAK1452 | mad10 deletion plasmid with Strep | pAK1127 | This study |
| pAK1384 | mad20 deletion plasmid with StrAB | pAK1127 | This study |
| pAK1268 | mad23 deletion with strAB for selection and upp counterselection (pCG116) | pAK1127 | This study |
| pAK1386 | mad25 deletion plasmid with StrAB | pAK1127 | This study |
| pAK1387 | mad26 deletion plasmid with StrAB | pAK1127 | This study |
| pAK1388 | mad28 deletion plasmid with StrAB | pAK1127 | This study |
| pAK966 | mamK deletion plasmid with strAB in pAK914 | pAK914 | This study |
| pAK1481 | pAK906–mad10 Complementation | pAK906 | This study |
| pAK1454 | pAK906–mad20 complementation | pAK906 | This study |
| pAK1507 | pAK906–mad23 complementation | pAK906 | This study |
| pAK1480 | pAK907–mad25 Complementation | pAK907 | This study |

| | | | |
|---|---|---|---|
| pAK1488 | pAK907–mad26 Complementation | pAK907 | This study |
| pAK1518 | pAK907– mad28 Complementation (with alt Start) | pAK907 | This study |
| pAK1489 | pAK906–mamK Complementation | pAK906 | This study |
| pAK318 | pKT25 | | EUROMEDEX |
| pAK319 | pKNT25 | | EUROMEDEX |
| pAK320 | pUT18 | | EUROMEDEX |
| pAK321 | pUT18C | | EUROMEDEX |
| pAK322 | pKT25-zip | | EUROMEDEX |
| pAK323 | pUT18C-zip | | EUROMEDEX |
| pAK1525 | KT25-mad10 | pAK318 | This study |
| pAK1526 | mad10-KNT25 | pAK319 | This study |
| pAK1527 | mad10-UT18 | pAK320 | This study |
| pAK1528 | UT18C-mad10 | pAK321 | This study |
| pAK1529 | KT25-mad20 | pAK318 | This study |
| pAK1530 | mad20-KNT25 | pAK319 | This study |
| pAK1531 | mad20-UT18 | pAK320 | This study |
| pAK1532 | UT18C-mad20 | pAK321 | This study |
| pAK1541 | KT25-mad23 | pAK318 | This study |
| pAK1542 | mad23-KNT25 | pAK319 | This study |
| pAK1543 | mad23-UT18 | pAK320 | This study |
| pAK1544 | UT18C-mad23 | pAK321 | This study |
| pAK1549 | KT25-mad25 | pAK318 | This study |
| pAK1550 | mad25-KNT25 | pAK319 | This study |
| pAK1551 | mad25-UT18 | pAK320 | This study |
| pAK1552 | UT18C-mad25 | pAK321 | This study |
| pAK1553 | KT25-mad26 | pAK318 | This study |
| pAK1554 | mad26-KNT25 | pAK319 | This study |
| pAK1555 | mad26-UT18 | pAK320 | This study |
| pAK1556 | UT18C-mad26 | pAK321 | This study |
| pAK1557 | KT25-mad28 | pAK318 | This study |
| pAK1558 | mad28-KNT25 | pAK319 | This study |
| pAK1559 | mad28-UT18 | pAK320 | This study |

| | | | |
|---|---|---|---|
| pAK1560 | UT18C-mad28 | pAK321 | This study |
| pAK1561 | KT25-mamK | pAK318 | This study |
| pAK1562 | mamK-KNT25 | pAK319 | This study |
| pAK1563 | mamK-UT18 | pAK320 | This study |
| pAK1564 | UT18C-mamK | pAK321 | This study |

**Supplemental Table 3.** *Table of all plasmids used in this study*

All of the plasmids used for this study and the previously published backbones used to generate the plasmids used for this research.

| Primer | Sequence | Description |
|---|---|---|
| VR123-F | AAACGCAAAAGAAAATGCCGATATCCTATTG GCCTCTAGAGGAATTCCAGGATTCCCTCA | mad20 deletion upstream |
| VR124-R | GGATCCCCCATCCACTAAATTTAAATAGGATC CTCTGTCCTTAATCTTGGCC | mad20 deletion upstream |
| VR125-F | GGATCCTATTTAAATTTAGTGGATGGGGGAT CCAGGATCGCCAACAATGCCTG | mad20 deletion downstream |
| VR126-R | TGGCGGTCATGGTCTTTTCGGCGAGCTTCTC TAGAAGCTCCGCTTCAAGAACCAC | mad20 deletion downstream |
| VR127-F | AAACGCAAAAGAAAATGCCGATATCCTATTG GCCTCTAGACAAGTGGAGTTAGGGTGATT | mad25 deletion upstream |
| VR128-R | CATATGCCCATCCACTAAATTTAAATACATAT GGTCAGCTTTAAAAAGGGCCA | mad25 deletion upstream |
| VR129-F | CATATGTATTTAAATTTAGTGGATGGGCATAT GTGTTCGGCCATTGCCGTTAT | mad25 deletion downstream |
| VR130-R | TGGTGGCGGTCATGGTCTTTTCGGCGAGCTT CTCTAGACCTCGTCTCTCTAACACAAA | mad25 deletion downstream |
| VR131-F | AAACGCAAAAGAAAATGCCGATATCCTATTG GCCTCTAGACATCAAAAGCATGGAAGGGC | mad26 deletion upstream |
| VR132-R | GGATCCCCCATCCACTAAATTTAAATAGGATC CTTCGCGCTGCATTCCGTCCA | mad26 deletion upstream |
| VR133-F | GGATCCTATTTAAATTTAGTGGATGGGGGAT CCACTATTGCCTTGAATAAATG | mad26 deletion downstream |

| | | |
|---|---|---|
| VR134-R | TGGTGGCGGTCATGGTCTTTTCGGCGAGCTT CTCTAGAGCATCCAAGCGTTCTTTTTG | mad26 deletion downstream |
| VR135-F | AAACGCAAAAGAAAATGCCGATATCCTATTG GCCTCTAGAAACGGTTTACAGCGGATACA | mad28 deletion upstream |
| VR136-R | GGATCCCCCATCCACTAAATTTAAATAGGATC CGCATCCAAGCGTTCTTTTTG | mad28 deletion upstream |
| VR137-F | GGATCCTATTTAAATTTAGTGGATGGGGGAT CCGGCCCTGACCGAGGCGGATT | mad28 deletion downstream |
| VR138-R | TGGCGGTCATGGTCTTTTCGGCGAGCTTCTC TAGACAAAATGGAGATTGGAGGAA | mad28 deletion downstream |
| VR147-F | AGGCGGGCATGGCCAAGATTAAGGACAGAG GATCCCTGCTAAAGGAAGCGGAACACG | Add StrAB-mad20-F |
| VR148-R | ATTCGCTATCAGGCATTGTTGGCGATCCTGG ATCCCTAGTATGACGTCTGTCGCACCT | Add StrAB-mad20-R |
| VR149-F | AGGGGCCATGGCCCTTTTTAAAGCTGACCAT ATGCTGCTAAAGGAAGCGGAACACG | Add StrAB-mad25-F |
| VR150-R | GAAAAAGCATAACGGCAATGGCCGAACACAT ATGCTAGTATGACGTCTGTCGCACCT | Add StrAB-mad25-R |
| VR151-F | GGGCAAACACTCCCGGTCGATTCCGGACAG GATCCCTGCTAAAGGAAGCGGAACACG | Add StrAB-mad26-F |
| VR152-R | ATCACAGTTCATTTATTCAAGGCAATAGTGGA TCCCTAGTATGACGTCTGTCGCACCT | Add StrAB-mad26-R |
| VR153-F | ATATTCGAGCAAAAAGAACGCTTGGATGCGG ATCCCTGCTAAAGGAAGCGGAACACG | Add StrAB-mad28-F |
| VR154-R | GGTATGGCTAATCCGCCTCGGTCAGGGCCG GATCCCTAGTATGACGTCTGTCGCACCT | Add StrAB-mad28-R |
| VR319-F | GTCATGGTCTTTTCGGCGAGCTTCTCTAGAA TCTCCAAAAAGTTGCTGGGC | mad10 deletion upstream |
| VR320-R | CCCATCCACTAAATTTAAATAGGATCCTCTTC CATGGCGTCCTCCG | mad10 deletion upstream |
| VR321-F | TATTTAAATTTAGTGGATGGGGGATCCCCAG TGCGCAGCCGGCCTGA | mad10 deletion downstream |

| | | |
|---|---|---|
| VR322-R | GAAAATGCCGATATCCTATTGGCCTCTAGAT CGCGGCCTGGGCC | mad10 deletion downstream |
| VR323-F | CGACAATGCGCGGAGGACGCCATGGAAGAG CTGCTAAAGGAAGCGGAACAC | Add StrAB-mad10-F |
| VR324-R | GAGATGATCTCAGGCCGGCTGCGCACTGGG CTAGTATGACGTCTGTCGCACC | Add StrAB-mad10-R |
| VR277-F | AGACAGGATGAGGATCGTTTCGCGTCGACTG ATCATCTCCCGGCGGG | mad20 complementation in pAK906 |
| VR278-R | GGGAATTCGAGCTCGGTACCCGGGGATCCT GCGCCGCTCCTGGATTC | mad20 complementation in pAK906 |
| CG220-R | GAGCTCGGTACCCGGGGATCCTCTAGAACTT GCTTTTCCGCCGTCATAC | mad23 complementation in pAK906 |
| CG221-F | CAAGAGACAGGATGAGGATCGTTTCGCGATG GAAGACGCCATGAGCTC | mad23 complementation in pAK906 |
| VR335-F | AAGCCAAGAAAAACGTCGCCAACGTCGACTA TGGCCCTTTTTAAAGCTGACG | mad25 complementation in pAK907 |
| VR336-R | GGGAATTCGAGCTCGGTACCCGGGGATCCT CTACCCGTCTGCGACGTC | mad25 complementation in pAK907 |
| VR337-F | AGACAGGATGAGGATCGTTTCGCGTCGACTA TGGAAGAAAATACCCGCTACA | mad10 complementation in pAK906 |
| VR338R | GGGAATTCGAGCTCGGTACCCGGGGATCCT TCAGGCCGGCTGCG | mad10 complementation in pAK906 |
| VR355-F | AAGCCAAGAAAAACGTCGCCAACGTCGACTA TGGACGGAATGCAGCGC | mad26 complementation in pAK907 |

| | | |
|---|---|---|
| VR356-R | GGGAATTCGAGCTCGGTACCCGGGGATCCTTCATTTATTCAAGGCAATAGTCAGT | mad26 complementation in pAK907 |
| VR357-F | AGACAGGATGAGGATCGTTTCGCGTCGACTATGTCCGGAAGCAACGTGC | mamK complementation in pAK906 |
| VR358-R | GGGAATTCGAGCTCGGTACCCGGGGATCCTTTAATCCTTTCGCAGCTCGTC | mamK complementation in pAK906 |
| VR375-F | AAGCCAAGAAAAACGTCGCCAACGTCGACTACAGCTCTGGCGCTCTTG | mad28 complementation in pAK907 (with pmad28 & alt-start) |
| VR372-R | GGGAATTCGAGCTCGGTACCCGGGGATCCTCTAATCCGCCTCGGTCAGG | mad28 complementation in pAK907 (with pmad28 & alt-start) |
| VR390-F | CGGGCTGCAGGGTCGACTCTAGAGGATCCCATGGAAGAAAATACCCGCTACA | KT25_mad10 |
| VR391-R | TCACGACGTTGTAAAACGACGGCCGAATTCTCAGGCCGGCTGCG | KT25_mad10 |
| VR392-F | ACAGCTATGACCATGATTACGCCAAGCTTGATGGAAGAAAATACCCGCTACA | KNT25_mad10-noStop |
| VR393-R | CCGGGGATCCTCTAGAGTCGACCTGCAGGCGGCCGGCTGCGCA | KNT25_mad10-noStop |
| VR394-F | ACAGCTATGACCATGATTACGCCAAGCTTGATGGAAGAAAATACCCGCTACA | UT18_mad10-NoStop |
| VR395-R | CCGGGGATCCTCTAGAGTCGACCTGCAGGCGGCCGGCTGCGCA | UT18_mad10-NoStop |
| VR396-F | CGCCACTGCAGGTCGACTCTAGAGGATCCCATGGAAGAAAATACCCGCTACA | UT18C_mad10 |
| VR397-R | TAGTTATATCGATGAATTCGAGCTCGGTACTCAGGCCGGCTGCG | UT18C_mad10 |
| VR398-F | CGGGCTGCAGGGTCGACTCTAGAGGATCCCATGGCCAAGATTAAGGACAGACT | KT25_mad20 |

| | | |
|---|---|---|
| VR399-R | TCACGACGTTGTAAAACGACGGCCGAATTCT CAGGCATTGTTGGCGATCC | KT25_mad20 |
| VR400-F | ACAGCTATGACCATGATTACGCCAAGCTTGA TGGCCAAGATTAAGGACAGACT | KNT25_mad20-noStop |
| VR401-R | CCGGGGATCCTCTAGAGTCGACCTGCAGGC GGCATTGTTGGCGATCCTCT | KNT25_mad20-noStop |
| VR402-F | ACAGCTATGACCATGATTACGCCAAGCTTGA TGGCCAAGATTAAGGACAGACT | UT18_mad20-NoStop |
| VR403-R | CCGGGGATCCTCTAGAGTCGACCTGCAGGC GGCATTGTTGGCGATCCTCT | UT18_mad20-NoStop |
| VR404-F | CGCCACTGCAGGTCGACTCTAGAGGATCCCA TGGCCAAGATTAAGGACAGACT | UT18C_mad20 |
| VR405-R | TAGTTATATCGATGAATTCGAGCTCGGTACTC AGGCATTGTTGGCGATCC | UT18C_mad20 |
| VR422-F | CGGGCTGCAGGGTCGACTCTAGAGGATCCC ATGGAAGACGCCATGAGCTC | KT25_mad23 |
| VR423-R | TCACGACGTTGTAAAACGACGGCCGAATTCC TAACTCCACTTGTAGCGCA | KT25_mad23 |
| VR424-F | ACAGCTATGACCATGATTACGCCAAGCTTGA TGGAAGACGCCATGAGCTC | KNT25_mad23-noStop |
| VR425-R | CCGGGGATCCTCTAGAGTCGACCTGCAGGC ACTCCACTTGTAGCGCATGC | KNT25_mad23-noStop |
| VR426-F | ACAGCTATGACCATGATTACGCCAAGCTTGA TGGAAGACGCCATGAGCTC | UT18_mad23-NoStop |
| VR427-R | CCGGGGATCCTCTAGAGTCGACCTGCAGGC ACTCCACTTGTAGCGCATGC | UT18_mad23-NoStop |
| VR428-F | CGCCACTGCAGGTCGACTCTAGAGGATCCCA TGGAAGACGCCATGAGCTC | UT18C_mad23 |
| VR429-R | TAGTTATATCGATGAATTCGAGCTCGGTACCT AACTCCACTTGTAGCGCA | UT18C_mad23 |
| VR438-F | CGGGCTGCAGGGTCGACTCTAGAGGATCCC ATGGCCCTTTTTAAAGCTGACG | KT25_mad25 |

| | | |
|---|---|---|
| VR439-R | TCACGACGTTGTAAAACGACGGCCGAATTCC TACCCGTCTGCGACGTC | KT25_mad25 |
| VR440-F | ACAGCTATGACCATGATTACGCCAAGCTTGA TGGCCCTTTTTAAAGCTGACG | KNT25_mad25-noStop |
| VR441-R | CCGGGGATCCTCTAGAGTCGACCTGCAGGC CCCGTCTGCGACGTCG | KNT25_mad25-noStop |
| VR442-F | ACAGCTATGACCATGATTACGCCAAGCTTGA TGGCCCTTTTTAAAGCTGACG | UT18_mad25-NoStop |
| VR443-R | CCGGGGATCCTCTAGAGTCGACCTGCAGGC CCCGTCTGCGACGTCG | UT18_mad25-NoStop |
| VR444-F | CGCCACTGCAGGTCGACTCTAGAGGATCCCA TGGCCCTTTTTAAAGCTGACG | UT18C_mad25 |
| VR445-R | TAGTTATATCGATGAATTCGAGCTCGGTACCT ACCCGTCTGCGACGTC | UT18C_mad25 |
| VR446-F | CGGGCTGCAGGGTCGACTCTAGAGGATCCC ATGGACGGAATGCAGCGC | KT25_mad26 |
| VR447-R | TCACGACGTTGTAAAACGACGGCCGAATTCT CATTTATTCAAGGCAATAGTCAGT | KT25_mad26 |
| VR448-F | ACAGCTATGACCATGATTACGCCAAGCTTGA TGGACGGAATGCAGCGC | KNT25_mad26-noStop |
| VR449-R | CCGGGGATCCTCTAGAGTCGACCTGCAGGC TTTATTCAAGGCAATAGTCAGTTCA | KNT25_mad26-noStop |
| VR450-F | ACAGCTATGACCATGATTACGCCAAGCTTGA TGGACGGAATGCAGCGC | UT18_mad26-NoStop |
| VR451-R | CCGGGGATCCTCTAGAGTCGACCTGCAGGC TTTATTCAAGGCAATAGTCAGTTCA | UT18_mad26-NoStop |
| VR452-F | CGCCACTGCAGGTCGACTCTAGAGGATCCCA TGGACGGAATGCAGCGC | UT18C_mad26 |
| VR453-R | TAGTTATATCGATGAATTCGAGCTCGGTACTC ATTTATTCAAGGCAATAGTCAGT | UT18C_mad26 |
| VR454-F | CGGGCTGCAGGGTCGACTCTAGAGGATCCC ATGTCCCAAGAGGATAAGGCG | KT25_mad28 |

| | | |
|---|---|---|
| VR455-R | TCACGACGTTGTAAAACGACGGCCGAATTCCTAATCCGCCTCGGTCAGG | KT25_mad28 |
| VR456-F | ACAGCTATGACCATGATTACGCCAAGCTTGATGTCCCAAGAGGATAAGGCG | KNT25_mad28-noStop |
| VR457-R | CCGGGGATCCTCTAGAGTCGACCTGCAGGCATCCGCCTCGGTCAGGG | KNT25_mad28-noStop |
| VR458-F | ACAGCTATGACCATGATTACGCCAAGCTTGATGTCCCAAGAGGATAAGGCG | UT18_mad28-NoStop |
| VR459-R | CCGGGGATCCTCTAGAGTCGACCTGCAGGCATCCGCCTCGGTCAGGG | UT18_mad28-NoStop |
| VR460-F | CGCCACTGCAGGTCGACTCTAGAGGATCCCATGTCCCAAGAGGATAAGGCG | UT18C_mad28 |
| VR461-R | TAGTTATATCGATGAATTCGAGCTCGGTACCTAATCCGCCTCGGTCAGG | UT18C_mad28 |
| VR462-F | CGGGCTGCAGGGTCGACTCTAGAGGATCCCATGTCCGGAAGCAACGTGC | KT25_mamK |
| VR463-R | TCACGACGTTGTAAAACGACGGCCGAATTCTTAATCCTTTCGCAGCTCGTC | KT25_mamK |
| VR464-F | ACAGCTATGACCATGATTACGCCAAGCTTGATGTCCGGAAGCAACGTGC | KNT25_mamK-noStop |
| VR465-R | CCGGGGATCCTCTAGAGTCGACCTGCAGGCATCCTTTCGCAGCTCGTCC | KNT25_mamK-noStop |
| VR466-F | ACAGCTATGACCATGATTACGCCAAGCTTGATGTCCGGAAGCAACGTGC | UT18_mamK-NoStop |
| VR467-R | CCGGGGATCCTCTAGAGTCGACCTGCAGGCATCCTTTCGCAGCTCGTCC | UT18_mamK-NoStop |
| VR468-F | CGCCACTGCAGGTCGACTCTAGAGGATCCCATGTCCGGAAGCAACGTGC | UT18C_mamK |
| VR469-R | TAGTTATATCGATGAATTCGAGCTCGGTACTTAATCCTTTCGCAGCTCGTC | UT18C_mamK |

**Supplemental Table 4.** *Table of all primers used in this study*

| | | **Compared with:** | **Mann-Whitney U test** | | **Welch's T-Test** | |
|---|---|---|---|---|---|---|
| **Figure** | **Strain/ Condition** | **Strain/ Condition** | **pValue** | **Significance** | **pValue** | **Significance** |
| Figure 3C | Early | Late | <0.0001 | **** | <0.0001 | **** |
| Figure 3D | Early | Late | <0.0001 | **** | <0.0001 | **** |
| Figure 3E | Early | Late | <0.0001 | **** | <0.0001 | **** |
| Figure 3I | Immature crystals | Mature crystals | --- | --- | 0.0006 | *** |

**Supplemental Table 5.** *Table of all statistical tests used for figure 3 in this study*

| | | **Compared with:** | **One way ANOVA test** | | **Mann-Whitney U test** | |
|---|---|---|---|---|---|---|
| **Figure** | **Strain/ Condition** | **Strain/ Condition** | **pValue** | **Significance** | **pValue** | **Significance** |
| Figure 4D | *fmpA* | WT | <0.0001 | **** | <0.0001 | **** |
| Figure 4D | *fmpB* | WT | <0.0001 | **** | <0.0001 | **** |
| Figure 4E | *fmpA* | WT | <0.0001 | **** | <0.0001 | **** |
| Figure 4E | *fmpB* | WT | <0.0001 | **** | <0.0001 | **** |
| Figure 4F | *fmpA* | WT | <0.0001 | **** | <0.0001 | **** |
| Figure 4F | *fmpB* | WT | <0.0001 | **** | <0.0001 | **** |

**Supplemental Table 6.** *Table of all statistical tests used for Figure 4 in this study*

| | | **Compared with:** | **One way ANOVA test** | | **Mann-Whitney U test** | |
|---|---|---|---|---|---|---|
| **Figure** | **Strain/ Condition** | **Strain/ Condition** | **pValue** | **Significance** | **pValue** | **Significance** |
| Figure 5N*i* | Δ*mamK* | WT | <0.0001 | **** | <0.0001 | **** |
| Figure 5N*i* | Δ*mad28* | WT | 0.9378 | ns | 0.4697 | ns |
| Figure 5N*ii* | Δ*mamK* | WT | <0.0001 | **** | <0.0001 | **** |

| | | | | | | |
|---|---|---|---|---|---|---|
| Figure 5N*ii* | Δ*mad28* | WT | 0.0703 | ns | 0.0053 | ** |

**Supplemental Table 7.** *Table of all statistical tests used for figure 5 in this study*

| | | Compared with: | One way ANOVA test | | Mann-Whitney U test | |
|---|---|---|---|---|---|---|
| **Supplemental Figure** | **Strain/ Condition** | **Strain/ Condition** | **pValue** | **Significance** | **pValue** | **Significance** |
| Figure S1D | Δ*mad10* | WT | 0.1364 | ns | 0.0005 | *** |
| Figure S1D | Δ*mad20* | WT | 0.7606 | ns | 0.3754 | ns |
| Figure S1D | Δ*mad23* | WT | 0.7606 | ns | 0.5328 | ns |
| Figure S1D | Δ*mad25* | WT | 0.3521 | ns | 0.0147 | * |
| Figure S1D | Δ*mad26* | WT | 0.4881 | ns | 0.111 | ns |
| Figure S1D | Δ*mad28* | WT | 0.7606 | ns | 0.3365 | ns |
| Figure S1D | Δ*mamK* | WT | <0.0001 | **** | <0.0001 | **** |

| | | Compared with: | One way ANOVA test | |
|---|---|---|---|---|
| **Supplemental Figure** | **Strain/Condition** | **Strain/Condition** | **pValue** | **Significance** |
| Figure S1E | Δ*mad10* | WT | <0.0001 | **** |
| Figure S1E | Δ*mad20* | WT | 0.0001 | *** |
| Figure S1E | Δ*mad23* | WT | 0.5127 | ns |
| Figure S1E | Δ*mad25* | WT | 0.9895 | ns |
| Figure S1E | Δ*mad26* | WT | 0.0953 | ns |
| Figure S1E | Δ*mad28* | WT | 0.0381 | * |
| Figure S1E | Δ*mamK* | WT | <0.0001 | **** |

| | | Compared with: | Kruskal-Wallis test | |
|---|---|---|---|---|
| **Supplemental Figure** | **Strain/Condition** | **Strain/Condition** | **pValue** | **Significance** |
| Figure S1F | Δ*mad10* | WT | <0.0001 | **** |
| Figure S1F | Δ*mad20* | WT | <0.0001 | **** |
| Figure S1F | Δ*mad23* | WT | 0.2744 | ns |

| | | | | |
|---|---|---|---|---|
| Figure S1F | Δ*mad25* | WT | 0.0073 | ** |
| Figure S1F | Δ*mad26* | WT | 0.0805 | ns |
| Figure S1F | Δ*mad28* | WT | 0.0007 | *** |
| Figure S1F | Δ*mamK* | WT | <0.0001 | **** |

**Supplemental Table 8.** *Tables of all statistical tests used for Supplemental Figure S1 in this study*

**References:**


1. Liu, Y., Van Den Ent, F. & Löwe, J. Filament structure and subcellular organization of the bacterial intermediate filament–like protein crescentin. *Proc. Natl. Acad. Sci.* **121**, e2309984121 (2024).
2. Pohl, A. *et al.* Decoding Biomineralization: Interaction of a Mad10-Derived Peptide with Magnetite Thin Films. *Nano Lett.* **19**, 8207–8215 (2019).
3. Paysan-Lafosse, T. *et al.* InterPro in 2022. *Nucleic Acids Res.* **51**, D418–D427 (2023).
4. Kearse, M. *et al.* Geneious Basic: An integrated and extendable desktop software platform for the organization and analysis of sequence data. *Bioinformatics* **28**, 1647–1649 (2012).
5. Hallgren, J. *et al.* DeepTMHMM predicts alpha and beta transmembrane proteins using deep neural networks. Preprint at https://doi.org/10.1101/2022.04.08.487609 (2022).
6. Uzun, M. *et al.* Recovery and genome reconstruction of novel magnetotactic Elusimicrobiota from bog soil. *ISME J.* **17**, 204–214 (2023).
7. Gilchrist, C. L. M. & Chooi, Y.-H. clinker & clustermap.js: automatic generation of gene cluster comparison figures. *Bioinformatics* **37**, 2473–2475 (2021).
8. Mirdita, M. *et al.* ColabFold: making protein folding accessible to all. *Nat. Methods* **19**, 679–682 (2022).
9. Meng, E. C. *et al.* UCSF ChimeraX: Tools for structure building and analysis. *Protein Sci. Publ. Protein Soc.* **32**, e4792 (2023).
10. Ludwiczak, J., Winski, A., Szczepaniak, K., Alva, V. & Dunin-Horkawicz, S. DeepCoil—a fast and accurate prediction of coiled-coil domains in protein sequences. *Bioinformatics* **35**, 2790–2795 (2019).

11. Li, J. & Pan, Y. Environmental Factors Affect Magnetite Magnetosome Synthesis in Magnetospirillum magneticum AMB-1: Implications for Biologically Controlled Mineralization. *Geomicrobiol. J.* **29**, 362–373 (2012).
12. Matsunaga, T. *et al.* Complete genome sequence of the facultative anaerobic magnetotactic bacterium Magnetospirillum sp. strain AMB-1. *DNA Res. Int. J. Rapid Publ. Rep. Genes Genomes* **12**, 157–166 (2005).
13. Maratea, D. & Blakemore, R. P. Aquaspirillum magnetotacticum sp. nov., a Magnetic Spirillum. *Int. J. Syst. Evol. Microbiol.* **31**, 452–455 (1981).
14. Smalley, M. D., Marinov, G. K., Bertani, L. E. & DeSalvo, G. Genome Sequence of Magnetospirillum magnetotacticum Strain MS-1. *Genome Announc.* **3**, e00233-15 (2015).
15. Dziuba, M. *et al.* Magnetospirillum caucaseum sp. nov., Magnetospirillum marisnigri sp. nov. and Magnetospirillum moscoviense sp. nov., freshwater magnetotactic bacteria isolated from three distinct geographical locations in European Russia. *Int. J. Syst. Evol. Microbiol.* **66**, 2069–2077 (2016).
16. Liu, P. *et al.* Key gene networks that control magnetosome biomineralization in magnetotactic bacteria. *Natl. Sci. Rev.* **10**, nwac238 (2023).
17. Monteil, C. L. *et al.* Repeated horizontal gene transfers triggered parallel evolution of magnetotaxis in two evolutionary divergent lineages of magnetotactic bacteria. *ISME J.* **14**, 1783–1794 (2020).
18. Ke, L. *et al.* Characteristics and optimised fermentation of a novel magnetotactic bacterium, Magnetospirillum sp. ME-1. *FEMS Microbiol. Lett.* **365**, fny052 (2018).

19. Ke, L., Liu, P., Liu, S. & Gao, M. Complete Genome Sequence of Magnetospirillum sp. ME-1, a Novel Magnetotactic Bacterium Isolated from East Lake, Wuhan, China. *Genome Announc.* **5**, e00485-17 (2017).

20. Wang, Y. *et al.* Characterizing and optimizing magnetosome production of Magnetospirillum sp. XM-1 isolated from Xi'an City Moat, China. *FEMS Microbiol. Lett.* **362**, fnv167 (2015).

21. Wang, Y. *et al.* Complete Genome Sequence of Magnetospirillum sp. Strain XM-1, Isolated from the Xi'an City Moat, China. *Genome Announc.* **4**, e01171-16 (2016).

22. Koziaeva, V. V. *et al.* Draft Genome Sequences of Two Magnetotactic Bacteria, Magnetospirillum moscoviense BB-1 and Magnetospirillum marisnigri SP-1. *Genome Announc.* **4**, e00814-16 (2016).

23. Koziaeva, V. V. *et al.* Magnetospirillum kuznetsovii sp. nov., a novel magnetotactic bacterium isolated from a lake in the Moscow region. *Int. J. Syst. Evol. Microbiol.* **69**, 1953–1959 (2019).

24. Fdez-Gubieda, M. L. *et al.* Magnetite Biomineralization in Magnetospirillum gryphiswaldense: Time-Resolved Magnetic and Structural Studies. *ACS Nano* **7**, 3297–3305 (2013).

25. Schleifer, K. H. *et al.* The Genus *Magnetospirillum* gen. nov. Description of *Magnetospirillum gryphiswaldense* sp. nov. and Transfer of *Aquaspirillum magnetotacticum* to *Magnetospirillum magnetotacticum* comb. nov. *Syst. Appl. Microbiol.* **14**, 379–385 (1991).

26. Wang, X. *et al.* Complete Genome Sequence of Magnetospirillum gryphiswaldense MSR-1. *Genome Announc.* **2**, e00171-14 (2014).

27. Lefèvre, C. T. *et al.* Insight into the Evolution of Magnetotaxis in Magnetospirillum spp., Based on *mam* Gene Phylogeny. *Appl. Environ. Microbiol.* **78**, 7238–7248 (2012).

28. Zhu, K. *et al.* Isolation and characterization of a marine magnetotactic spirillum axenic culture QH-2 from an intertidal zone of the China Sea. *Res. Microbiol.* **161**, 276–283 (2010).

29. Ji, B. *et al.* Comparative genomic analysis provides insights into the evolution and niche adaptation of marine Magnetospira sp. QH-2 strain. *Environ. Microbiol.* **16**, 525–544 (2014).

30. Monteil, C. L. *et al.* Genomic study of a novel magnetotactic Alphaproteobacteria uncovers the multiple ancestry of magnetotaxis. *Environ. Microbiol.* **20**, 4415–4430 (2018).

31. Bazylinski, D. A. *et al.* Magnetovibrio blakemorei gen. nov., sp. nov., a magnetotactic bacterium (Alphaproteobacteria: Rhodospirillaceae) isolated from a salt marsh. *Int. J. Syst. Evol. Microbiol.* **63**, 1824–1833 (2013).

32. Trubitsyn, D. *et al.* Draft Genome Sequence of Magnetovibrio blakemorei Strain MV-1, a Marine Vibrioid Magnetotactic Bacterium. *Genome Announc.* **4**, e01330-16 (2016).

33. Lefèvre, C. T. *et al.* Novel magnetite-producing magnetotactic bacteria belonging to the Gammaproteobacteria. *ISME J.* **6**, 440–450 (2012).

34. Geurink, C. *et al.* Complete Genome Sequence of Strain BW-2, a Magnetotactic Gammaproteobacterium in the Family Ectothiorhodospiraceae, Isolated from a Brackish Spring in Death Valley, California. *Microbiol. Resour. Announc.* **9**, e01144-19 (2020).

35. Li, J. *et al.* Single-Cell Resolution of Uncultured Magnetotactic Bacteria via Fluorescence-Coupled Electron Microscopy. *Appl. Environ. Microbiol.* **83**, e00409-17 (2017).

36. Trubitsyn, D. *et al.* Complete Genome Sequence of Strain SS-5, a Magnetotactic Gammaproteobacterium Isolated from the Salton Sea, a Shallow, Saline, Endorheic Rift

Lake Located on the San Andreas Fault in California. *Microbiol. Resour. Announc.* **10**, e00928-20 (2021).

37. Lefèvre, C. T., Bernadac, A., Yu-Zhang, K., Pradel, N. & Wu, L.-F. Isolation and characterization of a magnetotactic bacterial culture from the Mediterranean Sea. *Environ. Microbiol.* **11**, 1646–1657 (2009).
38. Ji, B. *et al.* The chimeric nature of the genomes of marine magnetotactic coccoid-ovoid bacteria defines a novel group of Proteobacteria. *Environ. Microbiol.* **19**, 1103–1119 (2017).
39. Meldrum, F. C., Mann, S., Heywood, B. R., Frankel, R. B. & Bazylinski, D. A. Electron microscopy study of magnetosomes in a cultured coccoid magnetotactic bacterium. *Proc. R. Soc. Lond. B Biol. Sci.* **251**, 231–236 (1993).
40. Schübbe, S. *et al.* Complete Genome Sequence of the Chemolithoautotrophic Marine Magnetotactic Coccus Strain MC-1. *Appl. Environ. Microbiol.* **75**, 4835–4852 (2009).
41. Morillo, V. *et al.* Isolation, cultivation and genomic analysis of magnetosome biomineralization genes of a new genus of South-seeking magnetotactic cocci within the Alphaproteobacteria. *Front. Microbiol.* **5**, (2014).
42. Koziaeva, V. *et al.* Genome-Based Metabolic Reconstruction of a Novel Uncultivated Freshwater Magnetotactic coccus "Ca. Magnetaquicoccus inordinatus" UR-1, and Proposal of a Candidate Family "Ca. Magnetaquicoccaceae". *Front. Microbiol.* **10**, (2019).
43. Abreu, F. *et al.* Cell Adhesion, Multicellular Morphology, and Magnetosome Distribution in the Multicellular Magnetotactic Prokaryote Candidatus Magnetoglobus multicellularis. *Microsc. Microanal.* **19**, 535–543 (2013).

44. Abreu, F. *et al.* ‘Candidatus Magnetoglobus multicellularis’, a multicellular, magnetotactic prokaryote from a hypersaline environment. *Int. J. Syst. Evol. Microbiol.* **57**, 1318–1322 (2007).

45. Abreu, F. *et al.* Deciphering unusual uncultured magnetotactic multicellular prokaryotes through genomics. *ISME J.* **8**, 1055–1068 (2014).

46. Leão, P. *et al.* Ultrastructure of ellipsoidal magnetotactic multicellular prokaryotes depicts their complex assemblage and cellular polarity in the context of magnetotaxis. *Environ. Microbiol.* **19**, 2151–2163 (2017).

47. Zhang, R. *et al.* Characterization and phylogenetic identification of a species of spherical multicellular magnetotactic prokaryotes that produces both magnetite and greigite crystals. *Res. Microbiol.* **165**, 481–489 (2014).

48. Kolinko, S., Richter, M., Glöckner, F.-O., Brachmann, A. & Schüler, D. Single-cell genomics reveals potential for magnetite and greigite biomineralization in an uncultivated multicellular magnetotactic prokaryote. *Environ. Microbiol. Rep.* **6**, 524–531 (2014).

49. Lefèvre, C. T. *et al.* A Cultured Greigite-Producing Magnetotactic Bacterium in a Novel Group of Sulfate-Reducing Bacteria. *Science* **334**, 1720–1723 (2011).

50. Lefèvre, C. T. *et al.* Comparative genomic analysis of magnetotactic bacteria from the Deltaproteobacteria provides new insights into magnetite and greigite magnetosome genes required for magnetotaxis. *Environ. Microbiol.* **15**, 2712–2735 (2013).

51. Sakaguchi, T., Burgess, J. G. & Matsunaga, T. Magnetite formation by a sulphate-reducing bacterium. *Nature* **365**, 47–49 (1993).

52. Sakaguchi, T., Arakaki, A. & Matsunaga, T. Desulfovibrio magneticus sp. nov., a novel sulfate-reducing bacterium that produces intracellular single-domain-sized magnetite particles. *Int. J. Syst. Evol. Microbiol.* **52**, 215–221 (2002).

53. Nakazawa, H. *et al.* Whole genome sequence of *Desulfovibrio magneticus* strain RS-1 revealed common gene clusters in magnetotactic bacteria. *Genome Res.* **19**, 1801–1808 (2009).

54. Shimoshige, H. *et al.* Isolation and cultivation of a novel sulfate-reducing magnetotactic bacterium belonging to the genus Desulfovibrio. *PLOS ONE* **16**, e0248313 (2021).

55. Lefèvre, C. T., Frankel, R. B., Pósfai, M., Prozorov, T. & Bazylinski, D. A. Isolation of obligately alkaliphilic magnetotactic bacteria from extremely alkaline environments. *Environ. Microbiol.* **13**, 2342–2350 (2011).

56. Li, J. *et al.* Intracellular silicification by early-branching magnetotactic bacteria. *Sci. Adv.* **8**, eabn6045 (2022).

57. Li, J. *et al.* Crystal growth of bullet-shaped magnetite in magnetotactic bacteria of the *Nitrospirae* phylum. *J. R. Soc. Interface* **12**, 20141288 (2015).

58. Lin, W. *et al.* Genomic insights into the uncultured genus 'Candidatus Magnetobacterium' in the phylum Nitrospirae. *ISME J.* **8**, 2463–2477 (2014).

59. Spring, S. *et al.* Dominating Role of an Unusual Magnetotactic Bacterium in the Microaerobic Zone of a Freshwater Sediment. *Appl. Environ. Microbiol.* **59**, 2397–2403 (1993).

60. Kolinko, S., Richter, M., Glöckner, F.-O., Brachmann, A. & Schüler, D. Single-cell genomics of uncultivated deep-branching magnetotactic bacteria reveals a conserved set of magnetosome genes. *Environ. Microbiol.* **18**, 21–37 (2016).

61. Uzun, M. *et al.* Detection of interphylum transfers of the magnetosome gene cluster in magnetotactic bacteria. *Front. Microbiol.* **13**, 945734 (2022).

62. Zhao, Y. *et al.* Insight into the metabolic potential and ecological function of a novel Magnetotactic Nitrospirota in coral reef habitat. *Front. Microbiol.* **14**, 1182330 (2023).

63. Kolinko, S. *et al.* Single-cell analysis reveals a novel uncultivated magnetotactic bacterium within the candidate division OP3. *Environ. Microbiol.* **14**, 1709–1721 (2012).

64. Rahn-Lee, L. *et al.* A Genetic Strategy for Probing the Functional Diversity of Magnetosome Formation. *PLoS Genet.* **11**, e1004811 (2015).

65. Grant, C. R., Rahn-lee, L. & Legault, K. N. Genome Editing Method for the Anaerobic Magnetotactic. *Appl. Environ. Microbiol.* **84**, 1–12 (2018).
